# Towards predictive ecological biogeography: predicting vegetation type using fine-scale measurements of the abiotic environment

**DOI:** 10.64898/2026.10.09.757380

**Authors:** Karl H. Palmquist, Sirio Belga Fedeli, Nicolas Lenner, Stanislas Leibler

## Abstract

One of the goals of ecological biogeography is to accurately predict where major vegetation types, such as forests or grasslands, can establish themselves. Alas, our current understanding of the mechanisms by which environmental variables jointly influence vegetation types does not allow for such predictions. Here, we compile a database of local measurements of soil, weather, topography, and bedrock lithology for 6404 sites across the continental United States, which we complement with information about the vegetation around the measurement sites. By applying machine learning methods to a subset of the sites, we are able to train a machine that predicts - using only information about abiotic variables - the presence of forests or grasslands with nearly 90% accuracy on the rest of the sites. We find that while weather variables are statistically important for these predictions, a few soil variables prove to be the most powerful predictors. By separating the combined contributions of weather and soil, we show that soil texture is a better predictor under dry conditions than under wet conditions, while pH is important predictor regardless of the weather. Furthermore, from 136 sites undergoing grassland-to-forest succession, our analysis indicates that soil pH changes with a certain lag relative to vegetation change. Sites that have recently undergone a transition exhibit a higher pH, closer to that of grasslands, despite the presence of trees. The pH then decreases by about one unit over a 30-year period of forest development. Ultimately, our results indicate that fine-scale environmental measurements and historical ecological data can serve as a basis for quantitative bio-geographical predictions, which could eventually guide targeted conservation interventions.

## Introduction

Anthropogenic land use has profoundly transformed the terrestrial biosphere, driving an estimated twofold reduction in global plant biomass since the dawn of civilization (1, 2). Conservation efforts seek to counteract this decline through a combination of targeted protection and species reintroduction (3). However, if these efforts fail to account for local ecological suitability, they risk not only high mortality rates among introduced species, but also the degradation of native habitats through ecologically mismatched plantings (4, 5). Ultimately, the success of interventions depends on predicting the types of vegetation, such as forests or grasslands, likely to establish themselves naturally on a given site.

The ability to make these predictions hinges on an ecological question: how do environmental variables jointly determine which vegetation types succeed in a given environment? This foundational question interested naturalists and scientists since antiquity: Theophrastus, the ancient Greek naturalist, detailed how the presence of specific vegetation types depends on abiotic variables, such as soil, sunlight, and moisture. During the 19th century, scientists built up the nascent field of plant biogeography by debating whether abiotic variables or biotic interactions were the primary drivers of geographic plant distribution. Focusing on biotic interactions, and influenced by the botanist Augustin de Candolle (6), Charles Darwin acknowledged that the broad physiological limits are set by climate, but argued that competition between plants is the primary driver dictating where specific species are located (7). Focusing instead on climatic abiotic variables, Alexander von Humboldt, and later researchers, relied on spatial averages of coarse temperature and precipitation measurements (sometimes across thousands of kilometers) to explain the geographic distribution of broad vegetation types (8–10).

Coarse spatial averaging can be useful for explaining the geographic distribution of major plant biomes. However, smoothing out soil and topographic variables, which vary over several meters, to obtain a coarse description at a scale of several kilometers results in a loss of potentially valuable information for predicting the presence of vegetation types at fine spatial scales. This limitation is particularly relevant for regions where topography or local soils appear to override the influence of regional climate, thus constituting exceptions to major vegetation trends. Examples include riparian forests that persist in otherwise treeless arid regions (11), and tropical grasslands that dominate very humid areas when located on localized clay soils (12). To explain such cases, early ecologists turned to fine-scale measurements of abiotic variables. Building on earlier work, Eugenius Warming’s 1895 study proposed that site-specific conditions, particularly soil properties and hydrology, directly govern the composition and organization of plant communities (13).

Despite Warming’s work, contemporary studies have largely followed Humboldt’s ideas. Few have sought to integrate fine-scale measurements into global predictions of plant distribution. Most theoretical models of global biomes assume that certain climatic boundaries constrain plant types and the thresholds associated with these boundaries are then adjusted to fit observed vegetation distributions (14). This approach cannot distinguish whether the models accurately capture how environmental variables determine vegetation, or whether they simply fit large-scale measurements to observations. By substituting large-scale spatial proxies for fine-scale measurements, theoretical models and modern machine learning approaches (15–17) share a common limitation: they smooth out soil and topography heterogeneity, thus concluding that these abiotic variables have only a secondary role in determining vegetation distributions.

What appears to be missing from these modern studies is an approach in the spirit of historical plant biogeography: coupling fine-scale measurements of environmental variables and vegetation across a vast geographic region to discover why plants grow in a given location. Such a study would incorporate variation across climates and landscapes using measurements at spatial scales that match those at which plants are growing. In this work, we attempt such an endeavor. First, we assemble a database of available fine-scale measurements of soil, weather, topography, and bedrock lithology variables across 6404 locations in the continental United States. These measurements are combined with direct observations of vegetation growing at the location, categorized as either forest or grassland. To these data, we apply explainable machine learning methods that enable us to both predict the vegetation type growing in a given location, and to decipher how environmental variables, such as soil pH and precipitation, jointly contribute to predictions. Furthermore, we provide an initial exploration into how more complex landscape attributes, such as recent ecological history, might influence the accuracy of such predictions.

## Results

### Dataset of abiotic environmental variables and vegetation type

We first assembled a dataset that enabled us to establish correlations between local abiotic environmental variables and local vegetation type. We regrouped abiotic environmental variables that differ substantially in both their measurement methodology and spatial resolution, reflecting the practical reality of how our source data variables, describing soil, weather, topography, and bedrock lithology, were collected. Soil data were drawn from pedon records that spanned nearly a century, from the late 1920’s to 2021 (18). We only used soil data that had reported geographic coordinates to five or more decimal places, corresponding to a positional precision on the order of 1 m. Soil variables included both chemical properties, such as pH and macronutrient concentrations, as well as physical texture. Because sand, silt, and clay fractions inherently sum to one, we parameterized soil texture using only two complementary variables to avoid mathematical co-linearity. The first is the sand fraction, which accounts for macro-porosity and rapid drainage. The second is a ratio for the fine-texture fraction of soil, 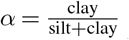, which captures the balance between water-retaining clay and infiltration-allowing silt. Long-term temporal weather averages were characterized by averaging both annual monthly precipitation and annual monthly temperature over 30 years, each estimated for the geographic position of individual pedon locations using a machine-learning model trained on weather station records across the continental United States. Because typical spacing between neighboring weather stations is on the order of 50 km, weather estimates reflect patterns at a much coarser spatial scale than the fine-scale measurements of soil. Topographic variables were derived from LiDAR-based elevation maps (19) at 10 m resolution. Bedrock lithology was assigned from the USGS State Geologic Map Compilation (SGMC) (20), which carries a spatial accuracy of approximately 500 m or better. Together, all these variables span nearly four orders of magnitude in spatial resolution: from meter-scale soil pedons to multi-kilometer-scale weather estimates, a mismatch that instigated the fine-to-coarse modeling framework developed here (Fig. 1).

**Fig. 1.**
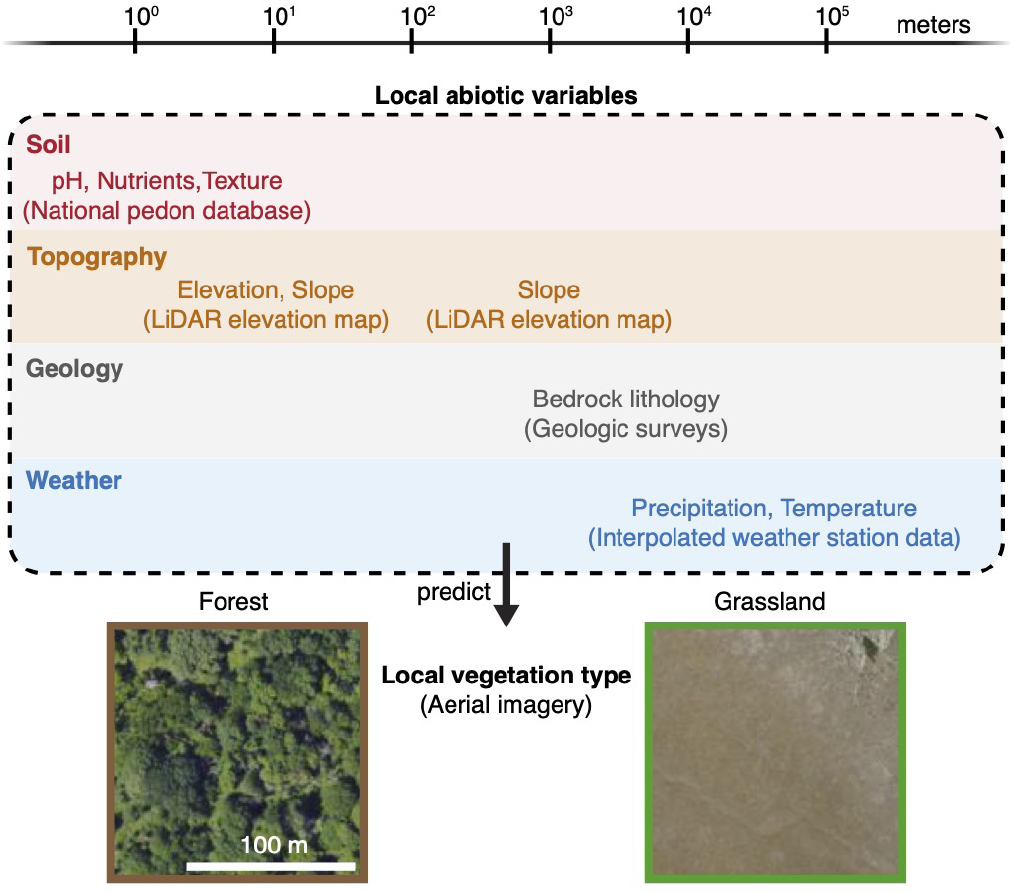
Schematic representation of dataset built to test if local abiotic environmental variables can predict local vegetation type. Soil measurements have a spatial resolution of 10^0^ m; topographic measurements have a horizontal measurement of 10^1^ m and are also averaged over 1 km; bedrock lithology maps have a spatial accuracy of at worst 500 m, and there is an average distance between weather stations on the order of 50 km.

Local vegetation type was determined through a two-step process. First, soil measurements were accompanied by categories of vegetation described by the field researchers at the time of soil collection. We filtered these descriptions retaining the samples that were labeled as “tree cover” or “grass/herbaceous cover.“ Second, we used the coordinates of the soil locations to extract 100 m × 100 m aerial images around each location. We retained only those locations for which, upon visual inspection, at least approximately 2*/*3 of the image area matched the database-assigned vegetation type label (Fig. 1). Although this double filtration process reduced the total number of locations to 6404 (from the total 9483), it increased our confidence in the assignment of the correct vegetation type at each location. In the following, for simplicity’s sake, we will refer to the two types simply as “forest“ and “grassland.“

### Classification of vegetation type from local abiotic environmental variables

To understand how the presence of a given vegetation type depends on abiotic environmental variables, it is instructive to first discuss their univariate distributions. When evaluating environmental variables independently—such as soil pH, precipitation, temperature, and soil texture (*α*)—there are distinct intervals along a given variable (*x*_*i*_) where the frequency distributions for forest and grassland are clearly distinct. Machine-learning models trained on these single variables achieve quite high classification accuracy within these intervals (typically located at the lowest or highest values of *x*_*i*_). However, the univariate distributions also exhibit significant regions of high overlap between forest and grassland sampling distributions. This inherent ambiguity indicates a need for a machine-learning approach that takes into account complex, non-linear multivariate environmental relationships when predicting the presence of a given vegetation type.

To disentangle these overlapping distributions and capture non-linear, multivariate environmental relationships, we trained an XGBoost classifier (21) on 80% of the 6404 sites. On remaining unused 20% test locations, the model achieved robust accuracies of 88.7 ± 0.2% for grasslands (class 0) and 89.3 ± 0.2% for forests (class 1), demonstrating a high level of generalization without systematic bias. Extensive cross-validation confirmed consistent performance regardless of the choice of training/test splits.

To determine how each of the 14 abiotic variables influenced predictions independently of the underlying algorithm, we calculated Shapley Additive exPlanations (SHAP) values *ϕ*_*i*_ (22). Because the prediction is expressed relative to a baseline dataset-average, the resulting SHAP values remain ecologically interpretable: *ϕ*_*i*_(*x*_*i*_) *>* 0 indicates a statistical association with forest cover, whereas *ϕ*_*i*_(*x*_*i*_) *<* 0 is associated with grassland. We then determined the statistical importance of variables by comparing the distribution of *ϕ*_*i*_(*x*_*i*_) for predictions using true vegetation-type labels against a null distribution generated using randomly permutated labels. We defined this null baseline as the 5th to 95th percentile range of the permuted *ϕ*_*i*_(*x*_*i*_) values. A variable was deemed significant if 60% or fewer of its spatial locations fell inside this random noise threshold.

Ranking the variables by 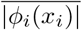 reveals that only four of the 14 predictors are statistically important relative to a null distribution (Fig. 2A). Soil pH emerged as the dominant predictor, followed by precipitation, soil texture (*α*), and temperature. Notably, a model restricted to soil variables outperformed one using only weather conditions (84.1 ± 0.1% vs. 76.5 ± 0.1% accuracy). While spatial weather variations are traditionally assumed to be the primary drivers of vegetation boundaries, these results indicate that fine-scale soil chemistry exerts a statistically greater association across our dataset.

**Fig. 2.**
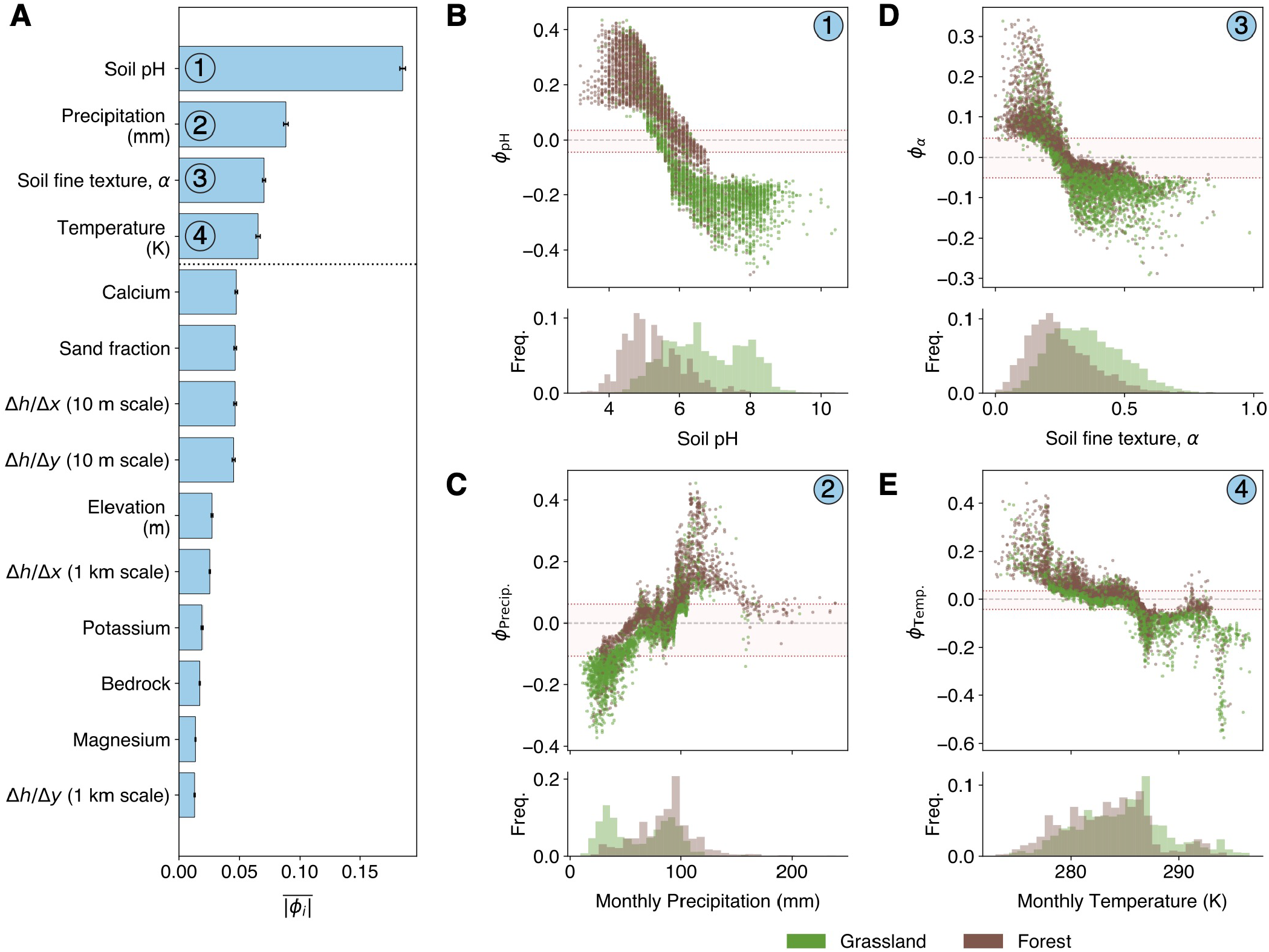
Importance of soil and environmental variables in determining vegetation type (grassland vs. forest) across the continental dataset. (A) Variables are ranked by their mean absolute value, 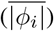, with error bars representing the SE of the mean. Variables above the dotted line are the four “major variables” discussed in the text, where 60% or fewer of the data points fall within the 5th to 95th percentile range of the null distribution of *ϕ*_*i*_ (red lines in B-E). (B-E) Importance values, *ϕ*_*i*_(*x*_*i*_), plotted as a function of the corresponding variable *x*_*i*_ for the four major variables (panels 1–4, corresponding to ranks in A): soil pH (H_2_ O), monthly precipitation (mm), *α* (Clay/Silt+Clay ratio), and temperature (K). In each panel, points are colored by vegetation type (green = grassland, brown = forest). Red dashed lines indicate significance thresholds used in (A). Significance is determined conservatively as the 5th to 95th percentile range of the null distribution of *ϕ*_*i*_, where *ϕ*_*i*_ is calculated from the original dataset where vegetation types are randomly shuffled. The distributions below each scatter plot show the relative frequency of grassland (green) and forest (brown) observations across the range of each variable. Numbers in circled labels correspond to variable ranks as shown in (A).

Examining the *ϕ*_*i*_ distributions clarifies this fact. The *x*_*i*_ intervals of greatest SHAP importance (Fig. 2B-E, upper panels) align with the intervals with the largest separations between univariate histograms (Fig. 2B-E, lower panels). Low pH and low *α* values (coarser, siltier soils) strongly predict the presence of forests (*ϕ*_*i*_ *>* 0), whereas high pH and high *α* values (finer, clayey soils) predict the presence of grasslands (*ϕ*_*i*_ *<* 0) (Fig. 2B,D). For precipitation and temperature, there are large ranges of values where *ϕ*_Precip_ and *ϕ*_Temp_ are non-significant (Fig. 2C,E). With that being said, drier and warmer regions show significantly negative *ϕ*_Precip_ and *ϕ*_Temp_, indicating that weather is statistically important for predictions of grassland under such weather conditions. Wetter and cooler regions, on the other hand, show significantly positive *ϕ*_Precip_ and *ϕ*_Temp_, indicating that such weather is statistically associated with predictions of forests (Fig. 2C,E).

### Predictive ability of soil texture depends on long-term weather averages

To explore the joint contributions of environmental variables, we investigated if soil’s influence on predictions is dependent on weather. We first asked if the contribution of the third-ranked variable—fine soil fraction (*α*)—is weather-dependent, a question inspired by Walter’s two-layer hypothesis (23–25). Walter proposed that trees and grasses share dry environments by using water at different depths: grasses rely on shallow moisture, while trees use deeper water. In drylands, trees can persist where soil is coarser and water infiltrates deeper. In regions where water is not limiting, texture becomes less relevant.

To test Walter’s hypothesis, we plotted forest and grassland samples on the graph spanned by the axes of precipitation and temperature (Fig. 3A). We defined three regions with different values of precipitation and temperature, ranging from cold and dry to warm and wet: an Arid (AR) region with low rainfall (25–50 mm/month; 277–281 K), a Sub-humid (SHU) transition zone (69–85 mm/month; 282–285 K), and a Humid (HU) region with high rainfall (90–100 mm/month; 286–288 K). To isolate soil effects from those linked to meteorological variations, we applied—prior to machine learning—an algorithm for sampling paired by weather conditions across the sites in each of these three regions. This approach rigorously controls for potential weather differences within each region, ensuring that the machine learning algorithm looks for correlations between vegetation type and abiotic variables for forests and grasslands that exist under the same weather conditions.

**Fig. 3.**
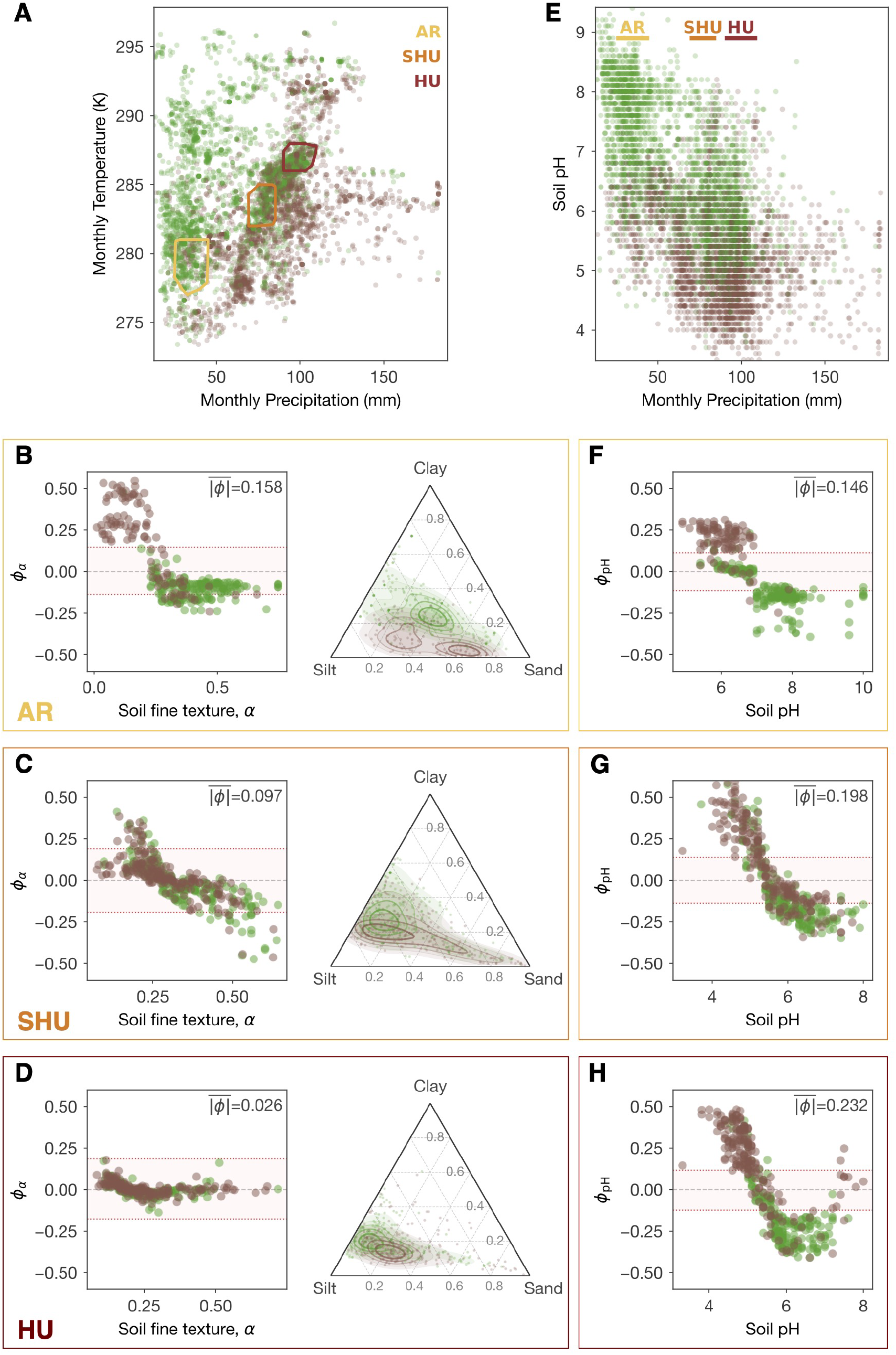
Importance of local soil variables across weather regimes. (A) Distribution of grassland and tree-cover samples in precipitation–temperature space. Dashed convex hulls delimit the three precipitation regimes: Arid (AR), Sub-humid (SH), and Humid (H). (B-D) SHAP dependence plots for soil texture (*α* = clay/(silt+clay)) across regimes, showing the per-sample contribution of *α* to vegetation classification. Positive values indicate a push toward tree cover; negative values toward grassland. Red dotted lines indicate the null permutation confidence threshold; 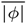 denotes mean absolute SHAP value. Center panels show the distribution of grassland and tree-cover samples in clay–silt–sand texture space, with kernel density contours overlaid for each vegetation class. (E) Precipitation versus soil pH for the full dataset; horizontal bars indicate the precipitation range of each matched regime. (F-H) SHAP dependence plots for soil pH across regimes, in the same format as B-D. Brown: forest; green: grassland.

In Fig. 3B-D, we compare the statistical association between vegetation type and texture (*α*) under dry and wet conditions. In dry environments, we observe a clear monotonically-decreasing trend between *ϕ*_*α*_ and *α*, indicating that low *α* values are statistically associated with predictions of forests, and high *α* values are statistically associated with predictions of grasslands (Fig. 3B). Under moderately wet (SHU) and wet (HU) conditions, this trend for *ϕ*_*α*_ becomes progressively weaker, indicating that *α* is no longer a statistically good predictor of the vegetation type (Fig. 3C,D). Soil texture triangles, where we plot sand, silt, and clay fractions for individual forests and grasslands, show a pattern that supports the machine-learning results. Under dry conditions, forests and grasslands are well separated, showing distinct cluster centers (Fig. 3B). However, under progressively wetter conditions (Fig. 3C,D), this separation between the clusters disappears, and the two vegetation types heavily overlap.

### Importance of soil pH does not depend on weather

Having established that the association between soil texture and vegetation cover depends on moisture, we now turn our attention to soil pH, which presents a particular challenge for interpretation. At the continental scale, soil pH co-varies strongly with precipitation (correlation coefficient *c*.*c*. = −0.62; Fig. 3E), a well-known phenomenon generally attributed to the balance between carbonate accumulation in arid regions and acidification through leaching under humid conditions (26, 27). This co-variation creates a potential confounding factor: the strong predictive signal of pH in global models (Fig. 2) could reflect either a direct relationship between pH and vegetation type or simply the simultaneous effect of precipitation on soil chemistry and vegetation cover.

To test whether this global association is merely an effect of precipitation, we evaluated the statistical relationship between soil pH and vegetation type within our weather-matched regions (Fig. 3A; AR, SHU, HU). Because these subsets rigorously control for weather differences, forest and grassland locations within each region receive statistically indistinguishable amounts of average rainfall. If the predictive power of pH were solely dependent on variations in precipitation, we would expect pH to lose its statistical importance within these controlled environments. Instead, the plots of *ϕ*_pH_ as a function of pH for all three regions reveal a consistent, monotonically decreasing trend (Fig. 3F–H). In each panel, *ϕ*_pH_ is positive at the lowest pH levels (statistically associated with forests) and drops below zero as pH increases (statistically associated with grasslands), with this transition consistently occurring between pH 5.5 and 6.0. This confirms that irrespective of long-term weather conditions, low pH is robustly associated with forests and high pH with grasslands.

### Soil chemistry changes as a function of ecological succession in vegetation type

Present-day ecosystems are shaped not only by current conditions but also by ecological and land-use legacies. History of land use can influence both current vegetation type and the environmental variables, including soil physicochemical properties, that are its best predictors. Studies occurring on individual sites have shown that forest succession can progressively alter soil characteristics (28, 29). However, it is largely unknown if the relationship between land use history and soil properties generalizes to different types of vegetation or weather and soil conditions. Using historical aerial imagery, we found a subset of 136 locations from our whole dataset for which we could track changes in land cover from grassland to forest. For these 136 locations, we identified the most recent year when the landscape was not forested (e.g., predominantly grassland). Using this year as our starting point, we created a time-alignment axis that describes the number of years that a site has subsequently been occupied by trees (i.e., the number of “years under forest“), with the end point being the year that soil was extracted from the location. This approach allowed us to assess how landscape history is associated with present vegetation and environmental conditions across an extended range of sites, thus establishing a link between past and present ecology.

We found that across the locations in this subset of our dataset, soil pH declines markedly with increasing years under forest, with median values falling from approximately 7.0 in the youngest stands (*<* 20 yr) to below 6.0 in stands older than 50 years (Fig. 4A,B). Most of the sites where the transition from grassland to forest was observed occur in moderate precipitation regimes (between on average 70 and 100 mm monthly). Within this range, pH values of younger forests (e.g., those that were more recently still non-forested) closely match those of grasslands. For older forests, we observed a transition toward lower pH values, close to the average values of all the forest population data (Fig. 4B).

**Fig. 4.**
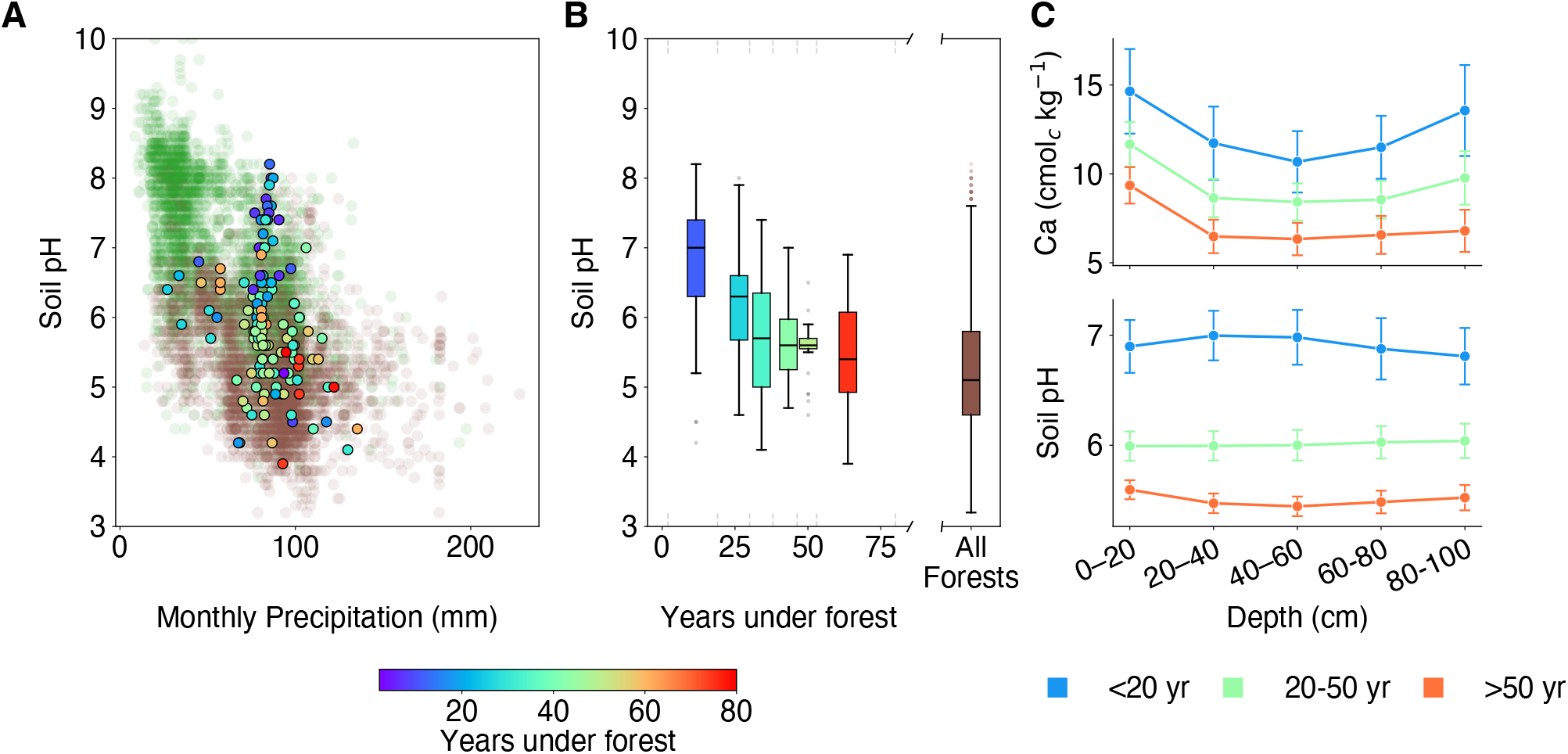
Changes in soil pH and exchangeable calcium as a function of years under forest. (A) Soil pH (H_2_ O) plotted against average monthly precipitation (mm) for the global grassland (green) and forest (brown) background dataset, with study sites overlaid as filled circles colored by years under forest (color scale below panel). (B) Box plots of soil pH (H_2_ O) for study sites grouped into six temporal quantiles, showing median, interquartile range, and 1.5*×* IQR whiskers, with outliers shown as grey points. Dashed vertical line segments provide the boundaries between each quantile. Brown boxplot on the right shows the distribution for all forest sites across the dataset. (C) Mean (*±* SE) exchangeable Ca (cmol_*c*_ kg^*−*1^ ; top) and soil pH (H_2_ O) (bottom) with soil depth (0–20, 20–40, 40–60, 60–80, and 80–100 cm) for the three age classes. Colors correspond to forest age: blue = *<*20 yr, green = 20–50 yr, orange = *>*50 yr.

Furthermore, the observed decrease in pH with forest age is relatively uniform with soil depth, up to 100 cm (Fig. 4C). In contrast, the concentration of exchangeable calcium decreases with soil depth, a trend that becomes more prominent as forest age increases (Fig. 4C). The depletion of calcium from deeper depths in the soil has previously been documented for mature forests (30). Our observation of a decrease in the concentration of exchangeable calcium as forests age is quantitative and statistically-significant support for the hypothesis that trees acidify soil as they grow by extracting base cations (e.g., calcium) from deeper soil layers (29).

## Discussion

### Overview

This study is rooted in the traditions of plant biogeography and macroecology—fields dedicated to understanding the geographic distribution of vegetation in relation to their surrounding environments (8, 14, 31–33). Our approach is to apply modern computational methods to explore a classic ecological question: why do certain types of vegetation, such as forest or grassland, grow where they do? We provide insight into this question by: first, establishing a database of fine-scale abiotic measurements across a large geographic region; and second, using machine-learning to analyze how well these measurements predict the presence of forest and grassland in nature.

### Predictive Accuracy and Generalizability

We demonstrate that the abiotic environment, such as soil conditions and weather, can predict the predominant vegetation type at 100 m scale across the continental United States (CONUS) with 89.0 ± 0.1% accuracy. We find that soil pH is statistically associated with vegetation type in a manner that is irrespective of long-term weather conditions. Moreover, our statistical learning machine demonstrates a high level of generalization to samples outside CONUS with classification accuracies of 62 ± 2% and 75 ± 2% for grassland and forest, respectively. This accuracy is achieved despite the fact that non-CONUS samples have fewer input variables available – only weather, soil texture, pH, and calcium concentration. However, there are particular non-CONUS regions where our predictions fail. For example, in the Uruguayan pampas, a region in South America mostly characterized by swaths of natural grasslands, our machine exhibits very low accuracy. This may be due to regional idiosyncrasies that are not captured by our input variables. For example, Darwin proposed the lack of trees in the pampas was due to excessive winds when he observed the landscape in 1839 (34), while more recent studies emphasize the role of grazing in modulating the presence of woody plants (35).

### Comparison with previous work

Recent machine learning studies have successfully used ensembles of decision trees or neural network approaches to predict global vegetation biome distributions (16, 17, 36, 37) and forest biomass across regions (38, 39), emphasizing climate and soil properties as key predictors at coarse spatial scales. Our study differs fundamentally in scale and data source: by integrating fine-scale, in-situ measurements of soil properties with local weather and topography, rather than relying on coarser remote sensing proxies or climate indices, we show that soil variables emerge as the strongest predictors of forest-grassland distribution when environmental heterogeneity is explicitly captured. This scale mismatch helps explain why prior studies privilege climate: coarser-resolution global models necessarily average over soil and topographic variation that dominates at local scales.

Additionally, our temporal analysis of grassland-to-forest succession reveals that soil pH lags behind vegetation shifts, declining approximately one unit over 30 years: a pattern consistent with vegetation-driven soil acidification documented in forest temporal sequences and long-term experiments (28, 29, 40). This legacy effect suggests that static prediction models, while useful for assessing current vegetation, must incorporate historical data for robust conservation guidance; a temporal dimension is clearly underexplored in existing frameworks.

### Limitations of the main results

Our results come with significant limitations. First, the variables we use and vegetation types we examine are limited by the spatial and temporal scales of their measurements. This ranges from a 1 m scale for soil properties values to an approximately 10^4^ m scale for weather conditions. Common approaches to increasing spatial resolution of environmental variables pose many challenges (41). Our approach, which uses raw weather station data within a relatively small geographic radius to predict 30-year precipitation and temperature averages, offers an alternative.

A future study or studies collecting measurements at increasing and decreasing coarseness could provide values for each variable over a range of length scales. Such a database could be then used to test the scale-dependence of environmental variables for predicting vegetation types.

Second, our approach implicitly assumes that abiotic variables and vegetation types are at steady state. We do not capture temporal environmental dynamics, such as short-term changes (e.g., seasonal shifts in soil chemistry) or longer-term changes (e.g., landscape or ecological history). Our initial results on ecological succession (Fig. 4) suggest that time dynamics will be important in future studies.

Finally, factors not included in this study, such as wildfires, pests or herbivorous grazing, also contribute to whether forest or grassland predominate over a given geographical region (15, 42–44). Future work should begin to assess how such factors statistically associate with vegetation type, and how they influence and complement the abiotic variables used in this study.

### Limitations for applied conservation

These limitations present specific challenges to conservation practitioners, who may be interested in using such works to inform targeted vegetation interventions. Specifically, practitioners need tools to help them decide where to intervene, whether the goal is to maintain a location’s current state or to restore its historical ecosystem. While we can successfully predict the current vegetation type, this is not the same as predicting what vegetation will successfully grow at a site in the future. This challenge arises because existing vegetation actively modifies its own soil. Take the example of a forest that was temporarily clear cut to establish grasslands for livestock pasture. If over time a young secondary forest reemerges, this area in transition might temporarily have a higher soil pH, because high pH is statistically associated with grassland, which existed recently. As a result, the machine might mistakenly classify the site as a grassland rather than a recovering forest in need of restoration and protection. Even if the location’s weather and soil texture are statistically associated with forest, this temporary pH signal can overpower the contributions of these other variables. That being said, we found that a machine trained using all of the inputs besides soil chemistry still has a predictive accuracy of 84.7 ± 0.1%. We can consider how incorporating a location’s ecological history as an input would address the issue of the landscape containing a memory of the past by allowing the machine to place less importance on transient soil chemistry in young stands. Until such historical variables can be integrated, conservation practitioners must couple these predictions with other information, such as historical site surveys.

An additional challenge arises because these machines learn strictly from past environmental conditions. Their reliability for guiding future conservation under changing climate remains an open question. To maintain forecast accuracy in the face of changing weather conditions, we must continue to collect fine-scale environmental data and consider incorporating temporal weather trajectories into our analyses, rather than limiting ourselves to simple long-term averages.

### Future directions

Ecological processes occur across space and time, presenting a challenge that requires more complete data across multiple scales (45–48). Future additional statistical data on abiotic and biotic factors should lead to a better understanding of the relative importance of abiotic and biotic conditions, and test hypotheses for how the presence and evolution of certain biota might influence abiotic variables. We hope that in the future, datasets will be extended to multiple spatiotemporal scales.

We expect that such multi-scale data will be particularly useful for predicting vegetation with finer phyto-geographic resolution, such as the presence of subclasses like conifers or hardwoods, or even of individual species. Recently, first attempts to predict the presence or absence of species have incorporated additional spatial information using convolutional neural networks (49).

Full multiscale analysis of such data, extending the one presented in this paper, may then enable studying how more mechanistic phenomena, occurring at smaller length or shorter time scales, influence larger-scale phenomena, such as the formation of patterns or behaviors visible only for large communities of plants.

## Methods

### Curation of Local Abiotic Environmental Conditions

The local abiotic environment was characterized by 14 variables obtained or derived from four independent sources. Soil data were drawn from the National Cooperative Soil Survey (NCSS)(18), which provides pedon-level soil profiles with standardized horizon descriptions and laboratory analyses; only NCSS records with geospatial precision of approximately 1 m were included. Raw soil variables were averaged over the top 30 cm of the soil horizon to produce pedon-level summary values. The resulting pedon dataset was merged with local weather averages estimated from nearby meteorological stations, topographic covariates derived from a 10 m LiDAR digital elevation model (elevation, slope) (19), and bedrock lithology assignments from the USGS State Geologic Map Compilation (20). Overall, this dataset contained 6404 samples. Using the geospatial coordinates of the soil pedons, samples were classified as forest or grassland when tree cover or grass/herbaceous cover, respectively, occupied more than two-thirds of the landscape (chosen to be ∼1 hectare square) based on satellite imagery (Google Maps) and matched the vegetation observed at the time of sampling (as recorded in the field).

### Assignment of Local Weather Estimates

To investigate localized historical climate dynamics, this study targeted the underlying, uncoarsened meteorological databases for precipitation and temperature managed by the University of East Anglia Climatic Research Unit (CRU) (50). Rather than utilizing spatially coarsened or interpolated grid models, individual point-source station observations spanning over 100 years were systematically mined directly from the original, quality-controlled (.clean.dtb) source database holdings to preserve native geographic variance and point-specific climate signals. While the core data collation mechanisms of these raw repositories follow the protocols described by Harris et al. (2020) (50), localized annual climate estimates representing a 30-year historical average prior to the exact date of soil sampling were generated for each pedon location using a custom XGBoost regressor framework. Local predictions were made by training models exclusively on raw weather stations within a strict focal radius of 300 km for precipitation and 500 km for temperature. These specific spatial bounds were empirically selected to minimize prediction error based on localized sensitivity analyses, corresponding to the radii that optimize cross-validated mean squared error and reflect the faster spatial decay of precipitation correlations relative to temperature. The localized regressors utilized latitude, longitude, elevation, and year as input variables to dynamically capture temporal and spatial variations across the sampling timelines. Model hyperparameters were optimized for each region via RandomizedSearchCV utilizing a 3-fold cross-validation scheme over 15 iterations. To ensure rigorous validation and counter spatial autocorrelation, optimization performance was evaluated against a strict spatial holdout of the nearest weather station (*R*^2^ = 0.96 for precipitation and *R*^2^ = 0.84 for temperature). The resulting localized estimates were assigned to each pedon and carried forward as variables in all subsequent analyses.

### Classifier and SHAP Decomposition

Vegetation type (forest vs. grassland) was treated as a binary outcome and predicted from the 14 local abiotic environmental conditions. Classification was performed with an XGBoost gradient-boosting classifier (xgboost v2.1.1). Hyperparameters— maximum tree depth, learning rate (*η*), and number of estimators—were optimized via GridSearchCV with 5-fold inner cross-validation. The full pipeline was repeated across five independent outer K-fold cross-validation loops, each initialized with a different random seed, to ensure that reported performance and variable importance estimates were not dependent on any particular data partition. We show the results of the model performance across inner crossvalidations and outer repeats.

SHAP values were computed for every test-fold observation using Kernel SHAP (KernelExplainer), applied to the hard classification output (0 = grassland, 1 = forest) rather than predicted probabilities, so that *ϕ*_*i*_ values remain directly interpretable in units of the class label. Per-sample SHAP vectors were stored across all five repetitions; the mean and standard deviation of *ϕ*_*i*_ were computed per sample across repetitions, and mean absolute SHAP values (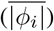) were then aggregated across all observations to produce a global ranking of variable importance.

### Null Variable-Importance Baseline

To establish a chance-level reference, vegetation-type labels were randomly permuted prior to each repetition of training—using a distinct random seed per repeat—and the complete modeling pipeline, including hyperparameter search, K-fold cross-validation, Kernel SHAP decomposition, and per-sample aggregation, was run identically on the shuffled labels. The resulting accuracy and SHAP distributions constitute the null baseline against which empirical variable importances were compared (Fig. 2A, dotted line).

### Construction of Climate-Matched Universes

To isolate the contribution of local soil variables from confounding variation in regional climate, sites were stratified into three discrete climate regimes—Arid (AR), Sub-humid (SHU), and Humid (HU)—defined by joint threshold on average monthly precipitation and average monthly temperature configured to bound specific regional clusters (AR: 25–45 mm, 277– 281 K; SHU: 69–85 mm, 282–285 K; HU: 90–110 mm, 286–288 K). Within each regime, a forest sample was drawn, and grassland sites were subsequently selected using nearest-neighbor matching across a dimensionless, *Z*-score standardized precipitation-temperature space to weight both climate dimensions evenly. This matching pipeline was executed with strict replacement constraints, ensuring that each grassland observation was uniquely assigned to at most one forest counterpart. This procedure constrains the forest and grassland populations within each regime to occupy statistically identical long-term precipitation and temperature distributions, such that any remaining difference in the abiotic profile reflects local soil or topographic conditions rather than regional macroclimate. The full matching and SHAP pipeline was repeated across five independent iterations (*N*_iterations_ = 5), each initialized with a distinct random seed, to ensure that downstream variable attributions were robust against alternative matching realizations.

### Successional History from Historical Aerial Imagery

From our pool of 6404 soil samples, we isolated 136 sites that were forested at the time of soil sampling, but for which grassland-to-forest succession was visually evident prior. We identified these successional transitions using historical aerial imagery from the Historic Aerials viewer (51). The most recent year in which a site appeared to be predominantly grassland (e.g., little-to-no tree cover visibly evident) was used to define the onset of forest establishment; the interval between that date and the year of soil sampling was recorded as years under forest. Changes in soil pH and exchangeable calcium concentration were then tracked as a function of years under forest, providing a temporal dimension to the cross-sectional patterns identified by the classifier.

## Data, Materials, and Software Availability

All data in this study were obtained from publicly available sources.

## ACKNOWLEDGEMENTS

The authors thank J. Chuang and S. Cohen for helpful discussions during the preparation of the manuscript. K.H.P was funded by the Martin and Helen Chooljian Fund in Biology and the Starr Foundation Fund in Biology. S.B.F. was funded by the Simons Foundation and the Charles L. Brown Fund in Biology. N.L. was supported by a stipend funded by the Simons Foundation, the Schmidt Fellow Fund, and the Eric and Wendy Schmidt Fund.

K.H.P., S.B.F., N.L., and S.L. designed research; K.H.P., N.L., and S.B.F. curated the main dataset, and K.H.P. performed research; K.H.P., N.L., S.B.F., and S.L. analyzed data; and K.H.P. and S.L. wrote the paper, with input from S.B.F. and N.L.

